# Changes in Myofibril Size, Shape, and Network Connectivity in Aging Muscle

**DOI:** 10.1101/2024.10.01.615981

**Authors:** Peter T. Ajayi, Zer Vue, Antentor O. Hinton, Brian Glancy

## Abstract

Across normal aging, striated muscles undergo structural remodeling associated with loss of force production. However, it is unknown how the organization of contractile myofibrillar networks, linked to their efficiency, is modified during aging. Using serial block-face scanning electron microscopy (SBF-SEM), we assess myofibril size, shape, and connectivity across different muscle types in young and geriatric mice and humans. Regardless of skeletal muscle fiber type in mice, age was associated with increased myofibrillar connectivity, with 24 months of age, as compared to 3 months, displaying more sarcomere branches. Distinctive age-related trends in myofibril size and shape were observed among each muscle type. Notably, there was a decrease in myofibril circularity from 3 months of age to 24 months of age in the gastrocnemius muscles of mice, contrasting with an increase in circularity in the soleus muscles during the same time frame. Additionally, while the soleus myofibrils in an aged cohort had a higher cross-sectional area, a reduction was observed in the gastrocnemius muscles. Cardiac muscles displayed no changes in sarcomere connectivity from 3 months to 24 months, although myofibril circularity and cross-sectional area were increased during this time. In human vastus lateralis muscles, sarcomere branching was positively correlated with advanced age. However, there were no consistent changes in myofibril size or shape across a wide age range from 16 to 68 years old. Overall, these data suggest that aging is associated with increased connectivity of the contractile networks within mammalian skeletal muscle.

**Graphical Abstract:** 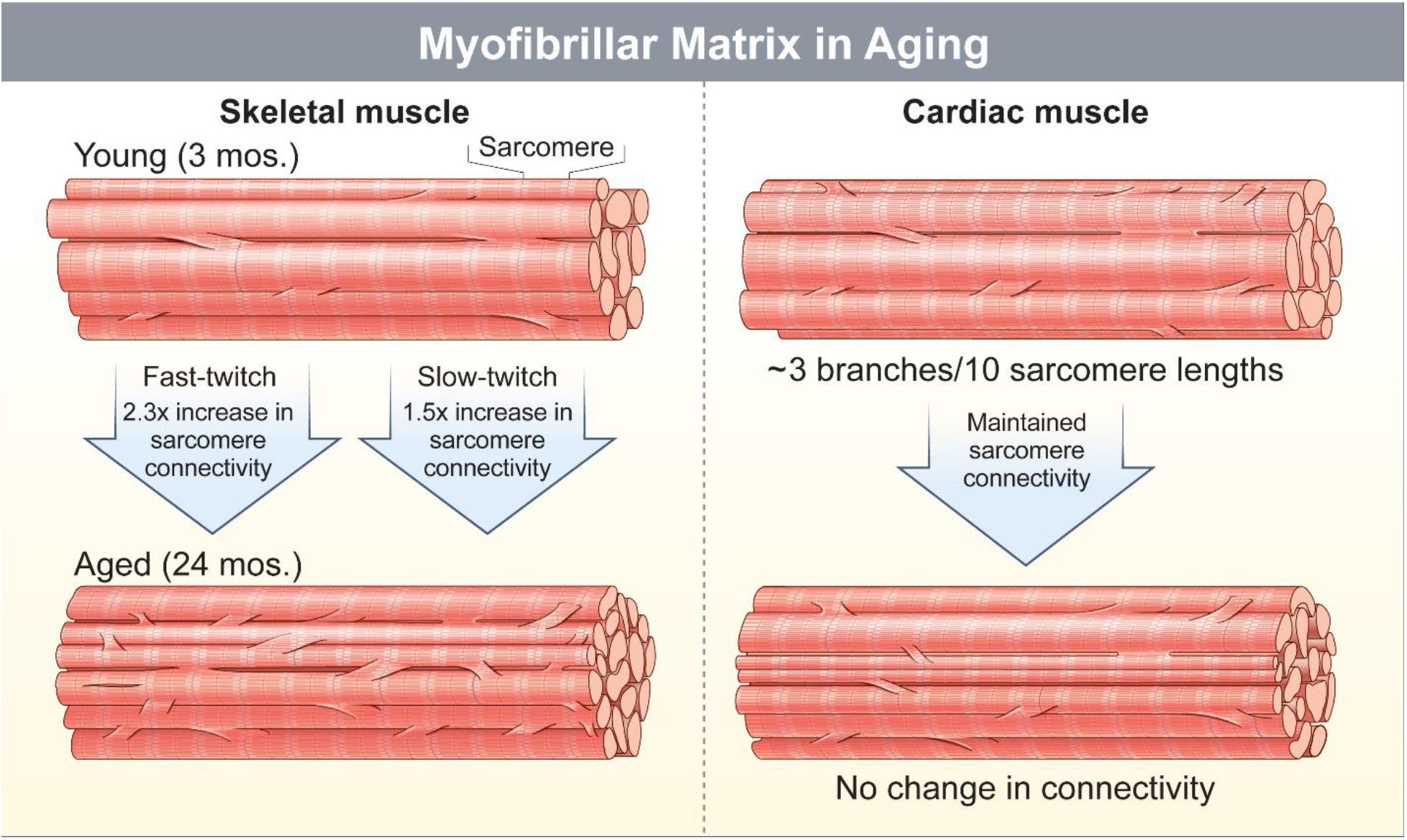

## Introduction

Aging is characterized by a progressive loss of muscle mass and strength termed sarcopenia (Yamada et al., 2022). Various interconnected factors are at play to induce sarcopenia, including muscle fiber atrophy (Dupont-Versteegden, 2005), apoptosis (Marzetti & Leeuwenburgh, 2006), reduced synthetic rates of proteins (Proctor, Balagopal, & Nair, 1998), hormonal changes (Kamel, Maas, & Duthie, 2002), chronic inflammation (Pan et al., 2021), mitochondrial dysfunction(Zer Vue et al., 2023), and impaired satellite cell function crucial for muscle regeneration (Alway, Myers, & Mohamed, 2014). The age-related decrease in muscle mass compromises strength and physical performance, contributing to challenges in maintaining overall balance and mobility (Visser et al., 2005). Sarcopenia imposes a significant economic burden in aging countries such as the US, with the total estimated cost of hospitalizations eclipsing $40 billion USD (Goates et al., 2019). Understanding the multifaceted nature of muscle loss with age is pivotal in developing effective strategies to enhance the quality of life for older individuals.

Sarcopenia is often accompanied by structural changes in muscle fibers, notably alterations in the size and number of sarcomeres, the functional unit of striated muscle cells(Lang et al., 2010). Structural changes in skeletal muscle are frequently reported first after the sixth decade in humans and followed by a progressive decline with increasing age (Oertel, 1986). Some reports suggest a preferential decrease in the size of fast-twitch fibers over slow-twitch fibers, leading to the preservation of muscle endurance at the expense of peak force (Coggan et al., 1992; L. Larsson, 1978). However, how this happens mechanistically at the ultrastructural level continues to be debated. Adjustments in sarcomere number and size have been proposed to mediate muscle atrophy (Sayed, Hibbert, Jorgenson, & Hornberger, 2023). Thinner myofibrils are frequently seen in aged muscles (Kaminska et al., 1998). The relationship between aging and sarcomere numbers, both in series and in parallel, remains limited; however, there is some evidence indicating a potential decrease (Hinks & Power, 2024). Sarcomere number is highly adjustable, being capable of either increasing or decreasing in adult muscles (Narici & Maganaris, 2007; Tabary, Tardieu, Tardieu, Tabary, & Gagnard, 1976; Williams & Goldspink, 1976). These described structural alterations have implications for overall muscle performance, impacting peak force and contraction speed during aging.

The bulk of the skeletal muscle volume is comprised of myofibrillar networks that work in tandem with the cytoskeleton to transmit longitudinal and lateral forces (Street, 1983; Willingham, Kim, Lindberg, Bleck, & Glancy, 2020). Branching of sarcomeres enables the formation of mesh-like myofibrillar networks. Sarcomere branching occurs through either a sarcomere split or a myofilament transfer mechanism. During a sarcomere split, one sarcomere transitions into two or more structures, which can be determined by the number of separated Z-discs. Structures usually occupying the splits’ fork include sarcotubular networks (sarcoplasmic reticulum + t-tubules) and mitochondria. Distinctively, myofilament transfers do not yield any additional sarcomeres as the myofilaments that are gained by one sarcomere are lost by the directly adjacent sarcomere. The frequency of sarcomere branching decreases during early-to-late postnatal development in mice and is more abundant in slow-twitch fibers compared to fast-twitch fibers (Willingham et al., 2020). These data suggest that the structural pathway for force transmission within a muscle cell is both dynamic over time and cell-type specific.

How, or whether, the muscle myofibrillar network is remodeled during aging remains unclear. Thus, we aimed to investigate how aging impacts the size, shape, and connectivity of sarcomeres in mammalian skeletal muscles and hypothesized that sarcomere connectivity would increase with age. To test this hypothesis, we evaluated muscle contractile structure using three-dimensional (3D) serial block face-scanning electron microscopy (SBF-SEM) images of cardiac, fast-, and slow-twitch muscles of young and old mice and skeletal muscles from humans aged 16 to 68.

## Methods

### Human subjects

Human specimens were collected at Vanderbilt University Medical Center and approved by the Vanderbilt University Institutional Review Board (IRB) under the title “Mitochondria in Aging and Disease --study of archived and autopsy tissue” with an associated IRB number of 231584. All cohort exclusions, namely existing conditions, are described previously (Zer Vue et al., 2023). Gross histology of patients are described in Table 1.

**Table 1.** Human Biopsy Characteristics.

| Age | Sex | History | Diagnosis | Histology |
| --- | --- | --- | --- | --- |
| 16 | F | Fatigue and elevated CK | No significant histopathologic abnormality | No abnormalities |
| 21 | F | Muscle weakness | No significant histopathologic abnormality | Mild atrophy |
| 29 | M | Elevated CK | No significant histopathologic abnormality | No abnormalities |
| 41 | M |  | No significant histopathologic abnormality | No abnormalities |
| 48 | F | Myopathy | No significant histopathologic abnormality | No abnormalities |
| 59 | M | Leg cramping and leg weakness | No significant histopathologic abnormality | Scattered mildly atrophic fibers |
| 68 | F | Proximal weakness and rash following flu shot. Mild inflammatory myopathy | Normal | No abnormalities |

### Animals

Animal care and euthanasia were performed as previously described(Lam et al., 2021). In accordance with protocols approved by the University of Iowa Animal Care and Use Committee (IACUC), all experiments were performed on male mice, C57Bl/6J. Animals were housed at 22 °C with a 12-h light, 12-h dark cycle with free access to water and standard chow.

### Serial block face scanning electron microscopy

Serial block face-scanning electron microscopy was performed, as previously described (Zer Vue et al., 2023). Briefly, isolated mice or human samples were cut into 1 mm3 samples, fixed with 2% glutaraldehyde in 0.1 M cacodylate buffer, and processed using a heavy metal protocol After tissue was treated with filtered 0.1% thiocarbohydrazide for 20 min and 2% osmium tetroxide for 30 min, tissues were incubated overnight in 1% uranyl acetate at 4°C. A 0.6% lead aspartate solution for 30 min at 60°C and graded acetone dilutions in dehydration were performed prior to embedding in fresh Epoxy TAAB 812 hard resin polymerization at 60°C for 36–48 hours. The block was sectioned for transmission electron microscopy (TEM) to identify the area of interest, trimmed to a 0.5 mm × 0.5 mm region of interest (ROI), and glued to an aluminum pin. Finally, the pin was placed into an FEI/Thermo Scientific Volumescope 2 scanning electron microscope imaging.

### Image segmentation

Myofibrillar segmentation was performed using a similar approach as in other works (Willingham et al., 2020). In brief, SBF-SEM image volumes with voxel sizes of 12×12×50 nm were adjusted to align the XY axes with the cross-section of the muscle cell. Segmentation utilized the interpolation feature in the TrakEM2 plugin of ImageJ to stitch together sequential tracings along a myofibrillar segment. The accuracy of the multi-color representation for the different myofibrillar segments was assessed by overlaying tracings with raw images.

### Quantitative analysis of myofibrillar networks

Using cross-sectional views of the muscle, sarcomeres in the first image were numbered, and their connectivity was tracked along the length of the muscle fiber through subsequent Z-discs. The number of times the sarcomeres split or merged, and the number of times myofilaments were transferred to or from adjacent sarcomeres in parallel were counted. Myofibrillar cross-sectional area (CSA) and circularity were measured by converting the traced structures to binary images and using the Analyze Particles plugin in ImageJ for each slice throughout the volume.

### Statistical Analysis

Using Prism 8 (GraphPad, San Diego, CA), we conducted all statistical analyses. Independent samples t-test was used to assess mean values between ages. A statistical significance level was set at P < 0.05.

### Data Availability

The raw SBF-SEM datasets for the present study have not been deposited in a public repository but are available from the corresponding author upon reasonable request.

## Results

### Contractile Connectivity in Mouse Fast-twitch Muscle Fibers

Contractile connectivity varies by muscle type (Ajayi et al., 2022; Willingham et al., 2020). Thus, to evaluate the effect of aging, we examined muscles predominantly composed of fast-twitch fibers (gastrocnemius), slow-twitch fibers (soleus), and cardiac muscles in mice. At 3 months of age, fast-twitch myofibrils in the gastrocnemius muscles had a greater cross-sectional area (2.1 ± 0.02 µm^2^) and higher circularity (0.66 ± 0.00) compared to those at 24 months of age (0.63 ± 0.00 µm^2^, 0.52 ± 0.00) (Fig. 1C). The frequency of sarcomere branches (Fig. 1D) was lowest at 3 months (2.9 ± 0.57 branches per 10 sarcomeres, *n* = 2 datasets, 2 cells, 60 myofibrils, 862 sarcomeres) than 24 months (7.3 ± 0.77 branches per 10 sarcomeres, *n* = 2 datasets, 2 cells, 95 myofibrils, 727 sarcomeres). Every myofibril assessed at 24 months of age had at least one branch. These data indicate that the increase in sarcomere branching between 3 and 24 months of age in fast-twitch muscles is accompanied by dynamic changes in size rather than shape to yield smaller myofibrils.

**Figure 1.**
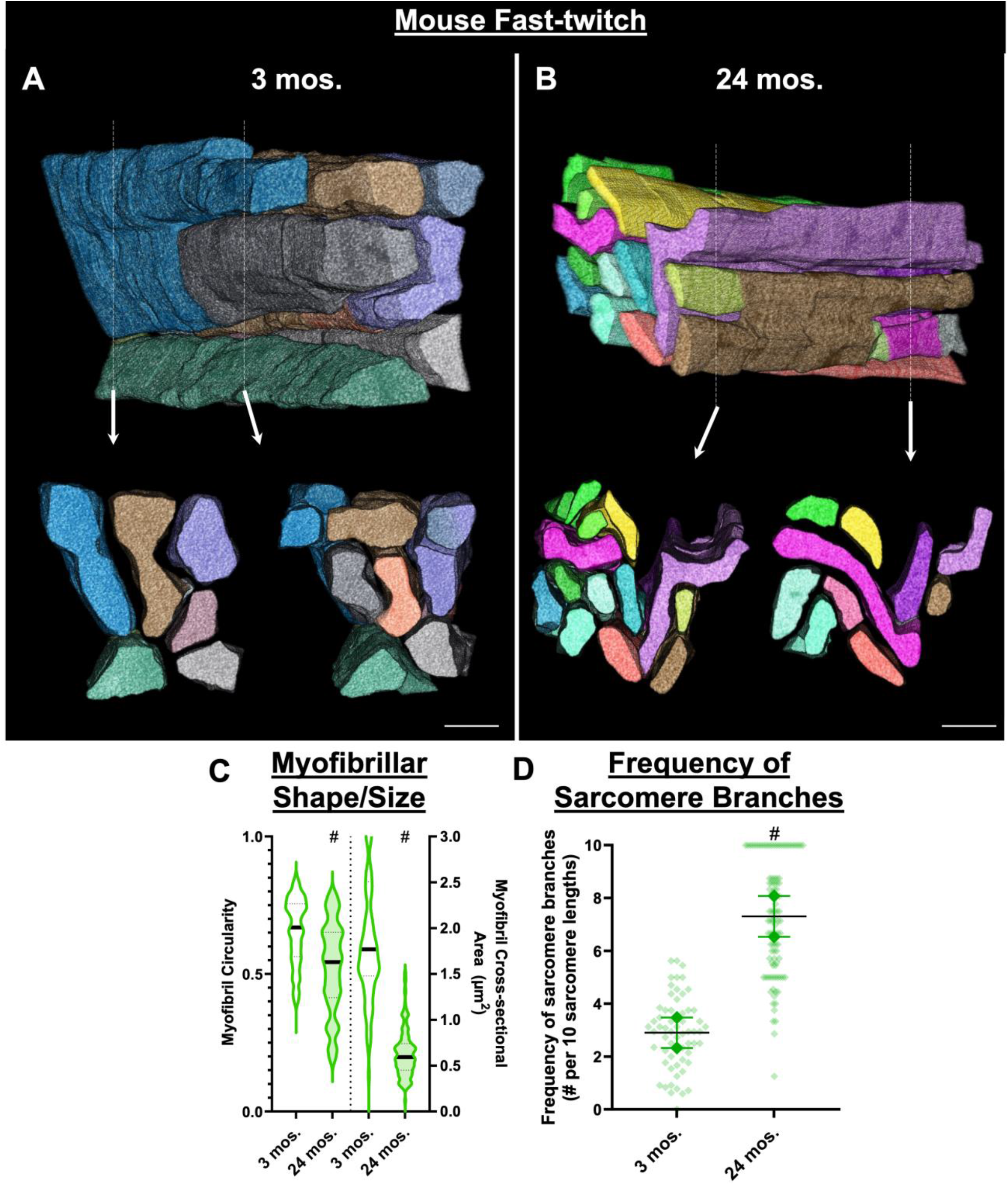
Myofibrillar networks in young and aged muscles in the gastrocnemius. **a, b** 3D rendering of myofibrillar networks at 3 and 24 months of age. Bottom two images show clippings through the long axis of the muscle directly above. Individual colors represent different myofibrillar segments separated by sarcomere branches. **c** Assessment of myofibril size and shape for 3 and 24 month aged muscles. **d** Frequency of sarcomere branching in 3 and 24 month old muscles. N values: 3 months—2 datasets, 2 cells, 60 myofibrils, 862 sarcomeres; and 24 months—2 datasets, 2 cells, 95 myofibrils,727 sarcomeres. Larger shape symbols represent data from a single dataset and smaller shape symbols represent data from a single myofibril. Bars represent muscle cell overall mean ± SE. Hash sign (#): Significantly different from 3 month old muscles (P < 0.01). Scale bars—2 μm.

### Contractile Connectivity in Mouse Slow-twitch Muscle Fibers

Slow-twitch muscle fibers, as characterized by their irregularly shaped sarcomeres, thicker z-discs, and higher mitochondrial content compared to fast-twitch fibers, display different trends in myofibrillar shape, size, and connectivity across age. At 24 months of age, myofibrils were more circular (0.61 ± 0.00) compared to myofibrils at 3 months of age (0.52 ± 0.00) and had a slightly greater cross-sectional area during the same period (0.72 ± 0.00 µm^2^, 0.66 ± 0.00 µm^2^) (Fig. 2C). The frequency of branches (Fig. 2D) unifying the myofibrillar matrix increased after 24 months (5.9 ± 0.36 branches per 10 sarcomeres, n = 3 datasets, 4 cells, 130 myofibrils, 1217 sarcomeres) from 3 months (4.0 ± 0.18 branches per 10 sarcomeres, n = 3 datasets, 5 cells, 95 myofibrils, 770 sarcomeres). All myofibrils assessed at both time points had at least one branch. The increase in sarcomere branching between 3 and 24 months is linked to more myofibrils with a circular profile and a slight increase in their sizes.

**Figure 2.**
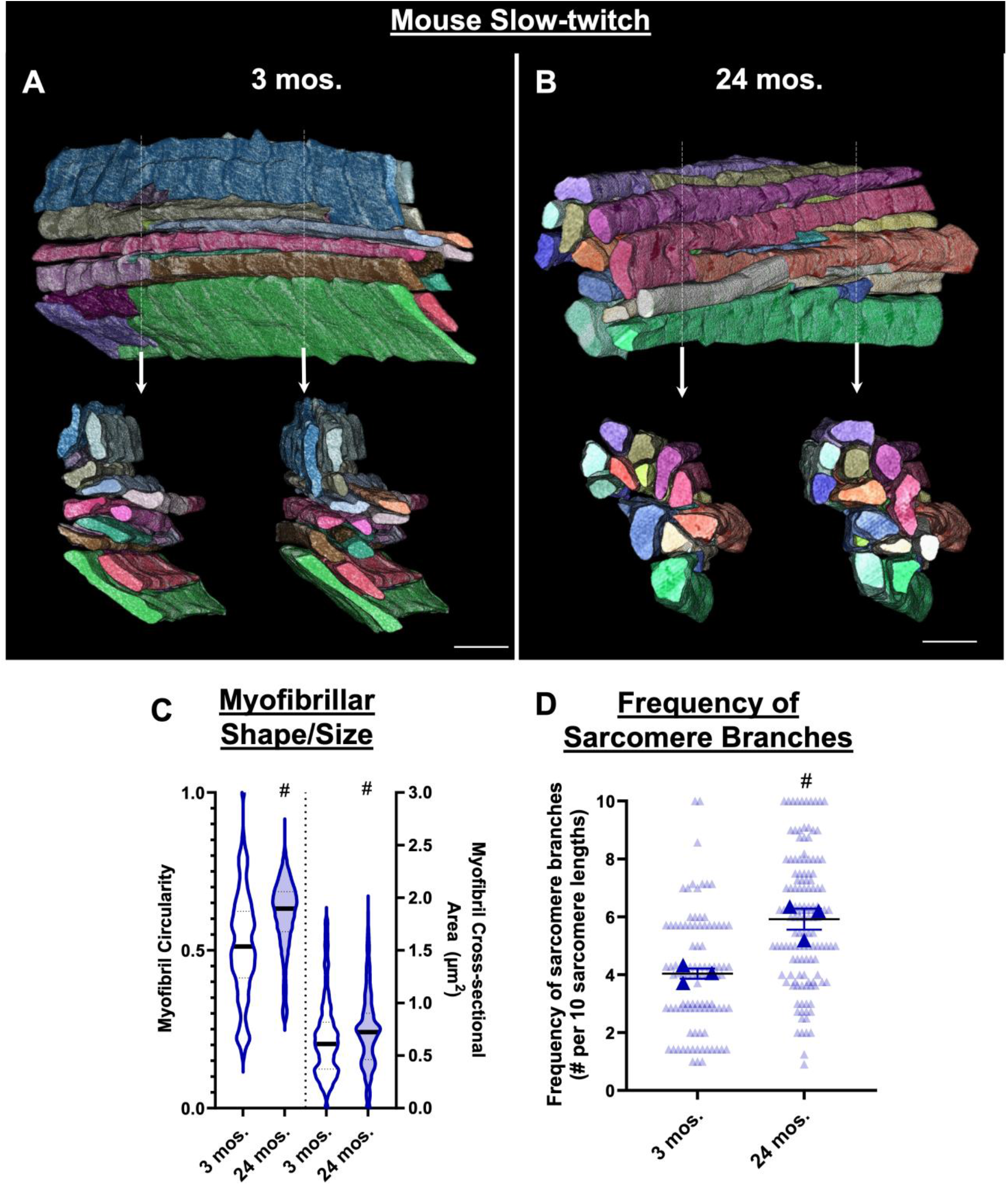
Myofibrillar networks in young and aged soleus muscles. **a, b** 3D rendering of myofibrillar networks at 3 and 24 months of age. Bottom two images show clippings through the long axis of the muscle directly above. Individual colors represent different myofibrillar segments separated by sarcomere branches. **c** Assessment of myofibril size and shape for 3 and 24 month aged muscles. **d** Frequency of sarcomere branching in 3 and 24 month old muscles. N values: 3 months—3 datasets, 4 cells, 95 myofibrils, 770 sarcomeres; and 24 months—3 datasets, 5 cells, 130 myofibrils, 1217 sarcomeres. Larger shape symbols represent data from a single dataset and smaller shape symbols represent data from a single myofibril. Bars represent muscle cell overall mean ± SE. Hash sign (#): Significantly different from 3 month old muscles (P < 0.01). Scale bars—2 μm.

### Contractile Connectivity in Mouse Cardiac Muscle Fibers

To determine whether the age-dependent changes in branching frequency were limited to skeletal muscle, we next evaluated the contractile structures from cardiac muscle cells. Unlike skeletal muscle cells, which contract voluntarily for skeletal movement and posture, cardiac muscle cells in the heart are under involuntary control for the sustained contractions required to pump blood through the body. Cardiac myofibrils display a greater circular profile (0.60 ± 0.00) and cross-sectional area (1.1 ± 0.01 µm^2^) at 24 months than myofibrils at 3 months of age (0.56 ± 0.00, 0.93 ± 0.01 µm^2^) (Fig. 3C). Additionally, the number of branches unifying the matrix remained unchanged during this time with 2.7 ± 0.49 branches per 10 sarcomeres at 3 months (n = 2 cells, 50 myofibrils, 649 sarcomeres) and 2.8 ± 0.32 branches per 10 sarcomeres at 24 months (n = 4 cells, 80 myofibrils, 800 sarcomeres) (Fig. 3D). Sarcomere branching appears to be stable across normal aging in mouse cardiomyocytes.

**Figure 3.**
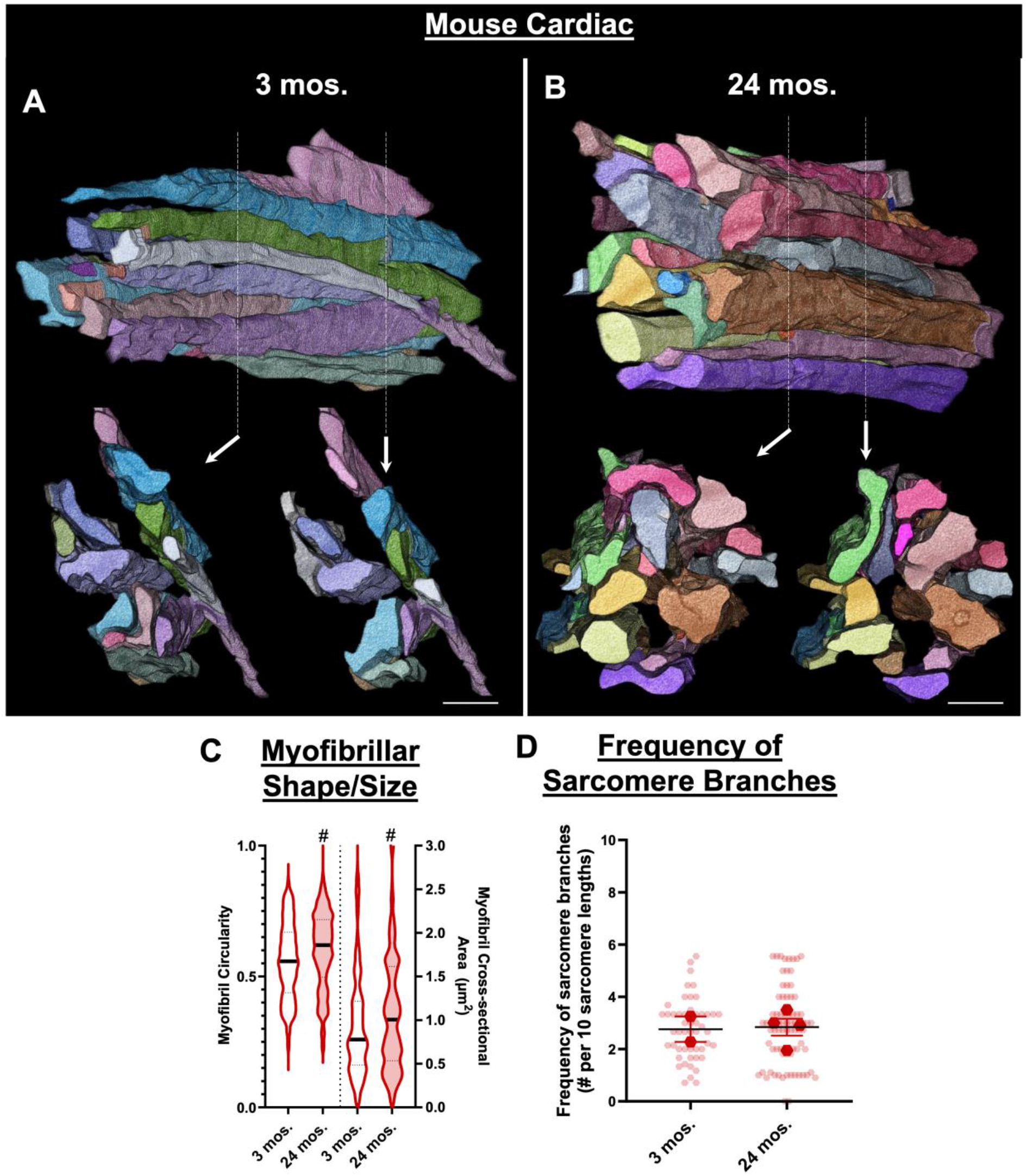
Myofibrillar networks in young and aged cardiac muscles.**a, b** 3D rendering of myofibrillar networks at 3 and 24 months of age. Bottom two images show clippings through the long axis of the muscle directly above. Individual colors represent different myofibrillar segments separated by sarcomere branches. **c** Assessment of myofibril size and shape for 3 and 24mmonth aged muscles. **d** Frequency of sarcomere branching in 3 and 24 month old muscles. N values: 3 months—2 cells, 50 myofibrils, 649 sarcomeres; and 24 months—2 cells, 80 myofibrils, 800 sarcomeres. Larger shape symbols represent data from a single cell and smaller shape symbols represent data from a single myofibril. Bars represent muscle cell overall mean ± SE. Hash sign (#): Significantly different from 3 month old muscles (P < 0.01). Scale bars—2 μm.

### Contractile Connectivity in Human Vastus Lateralis Muscle Fibers

To further evaluate whether sarcomere branching is dynamic across age in skeletal muscle, we assessed myofibrillar structures of the fast-twitch vastus lateralis of seven humans from different age groups (Table 1; Fig. 4 A-D). From ages 16 to 29, the myofibrils exhibited an increasingly circular appearance, but a significant decline was observed after the age of 29, with a notable increase at 68 years (Fig. 4E). Myofibril cross-sectional area was variable across age (Fig. 4F). We then compared the magnitude of sarcomere branching versus the age of the muscle sample for all seven humans which revealed a linear relationship (Fig. 4G,*R*^*2*^ = 0.5811, *p* = 0.0463) indicating that sarcomere branching increases with normal aging, which has important implications for understanding muscle function and developing strategies to address age-related muscle decline.

**Figure 4.**
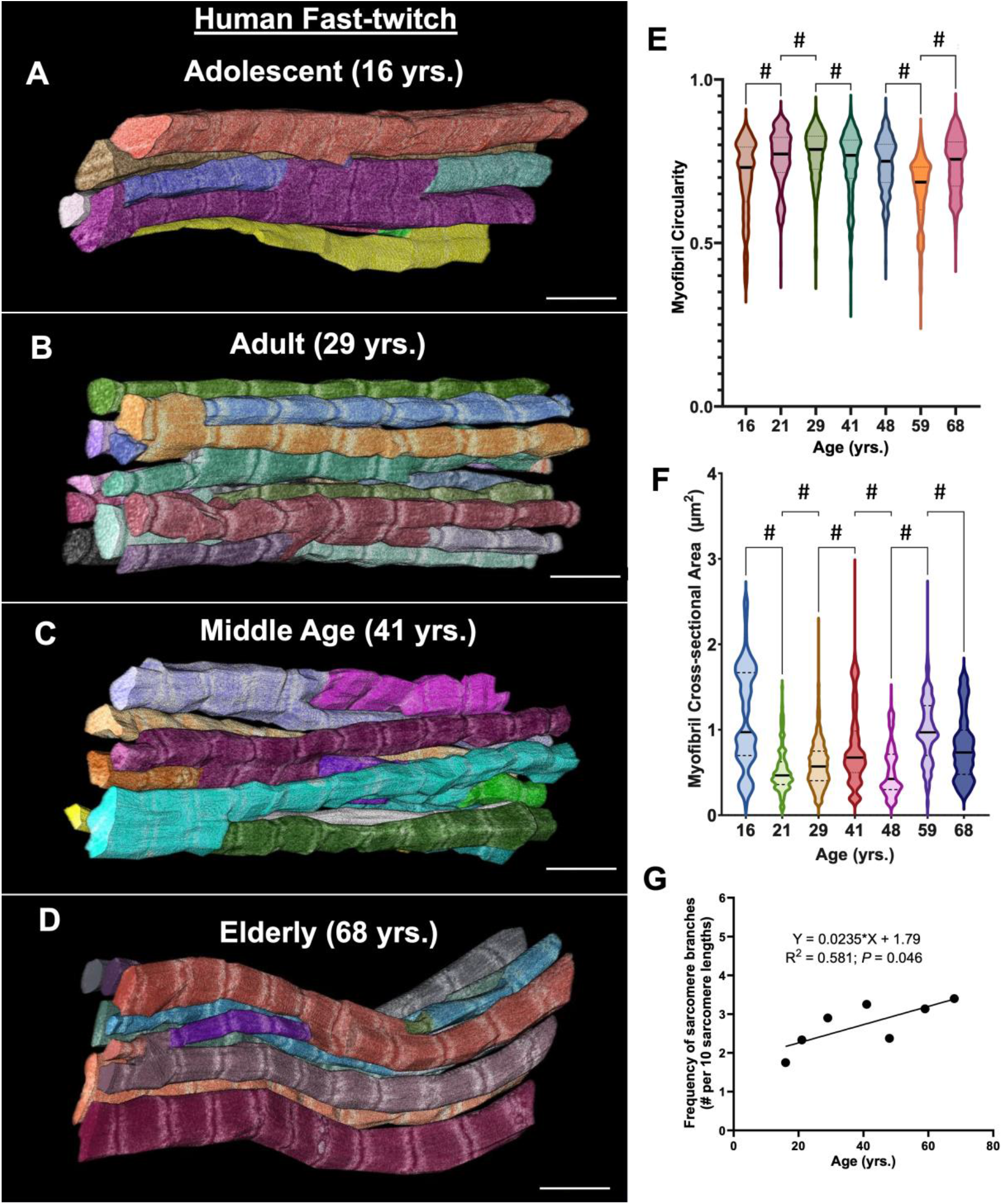
Myofibrillar networks in adolescent, adult, middle age, and elderly muscles. **a-d** 3D rendering of myofibrillar matrix across different age groups. Individual colors represent different myofibrillar segments separated by sarcomere branches. **E, f** Assessment of myofibril shape and size for muscles across the lifespan (16-68 years of age). **G** Frequency of sarcomere branching across age. N values: 16 years— 20 myofibrils, 200 sarcomeres; 21 years— 20 myofibrils, 180 sarcomeres; 29 years— 20 myofibrils, 200 sarcomeres; 41 years— 20 myofibrils, 200 sarcomeres; 48 years— 20 myofibrils, 139 sarcomeres; 59 years— 18 myofibrils, 252 sarcomeres; 68 years— 20 myofibrils, 100 sarcomeres. Circle symbols represent composite data of myofibrils from a single cell. Hash sign (#): Significantly different from prior muscles at previous age (P < 0.01). Scale bars—2 μm.

## Discussion

Here we show that the number of branching sarcomeres that unify myofibrillar networks increases with age in mammalian skeletal muscles (Figs. 1,2,4). Aligned with prior works (Ajayi et al., 2022; Willingham et al., 2020) the degree of connectivity within the contractile apparatus is muscle type dependent with more oxidative and lower force-producing muscles containing more branches. The change in magnitude across age is also fiber type-dependent with fast-twitch and slow-twitch fibers having 151.72% and 47.50% more branching after 24 months compared to 3 months, respectively (Fig. 1D and Fig. 2D). Despite initially having higher branch numbers, slow-twitch fibers incorporate less branching over time relative to fast-twitch fibers. Our comparison of the size, shape, and network connectivity of the 24-month-old myofibrils in both fiber types revealed similarities. This includes having similar sizes (CSA mean difference = 0.09 ± 0.01; P < 0.01), along with similar branching frequencies (sarcomere branching mean difference = 1.38 ± 0.74; P > 0.05). Thus, fast-twitch fibers appear to undergo more significant alterations in their myofibrillar architecture over time and exhibit characteristics more akin to slow-twitch fibers. These data are consistent with the fast-to slow-twitch transitions in motor units and myosin isoforms shown previously(Lars Larsson et al., 2019). While we did not observe consistent changes in the shape and size of myofibrils across different ages in human fast-twitch muscles, a positive correlation between age and branching frequency was still appreciated (Fig. 4G). We extend findings from previous work showing that fast-twitch muscles are preferentially downsized with age, whereas slow-twitch fibers are more or less preserved with age (L. Larsson, 1978).

Previously, it was hypothesized that muscle growth was a primary driver of sarcomere branching (Goldspink, 1970). However, between 3 months and 24 months as per the timeline in this present study, there is very little growth happening (Lin et al., 2018). Furthermore, when we pooled the data across fiber types, older muscles had 65 ± 4.6 percent of their sarcomeres occupied by branches compared to the 36 ± 3.5 percent seen in younger muscles (P < 0.01). Here, branching appears to be related to the loss of sarcomeres, as would be expected in sarcopenia (Thom, Morse, Birch, & Narici, 2007), rather than the increase in number associated with muscle growth (Jorgenson et al., 2024).

The increased branching, and hence network connectivity, in older muscles, may suggest a need to reroute forces around damaged regions of the muscle. Z-disc disruptions are a well-characterized feature of myofibrillar damage/remodeling (Yu, Carlsson, & Thornell, 2004). Based on single 2D images, it was previously shown that about 2% of fibers from healthy young people contain evidence of Z-disc streaming (Meltzer, Kuncl, & Yang, 1976). For a direct comparison of muscles from healthy younger (20-30-year-old) and elderly (65-75-year-old) men and women revealed similar percentages of muscle damage at around 3% (Roth et al., 2000). Consistently, our analysis of complete 3D structures of over 10,000 sarcomeres also showed little sarcomere disorganization as Z-discs were properly aligned in parallel and I-bands were present in the selected fields of view from young and aged muscles of both mice and humans. Thus, the observed increase in branching in older skeletal muscles here does not appear to be directly related to sarcomere damage.

In a more metabolic-intensive muscle type such as cardiomyocytes, contractile connectivity remains stable from young adult to old age in mice (Fig. 3D). We previously showed that the percentage of sarcomere branching is highly variable across mouse cardiac postnatal development with the percentage of branches decreasing by over 50% from birth to postnatal day 42, resulting in roughly 3 branches per 10 sarcomere lengths (Kim, Ajayi, Bleck, & Glancy, 2022). During this time, cardiomyocytes have already exited the cell cycle and shifted from proliferative growth to hypertrophic growth, where the cells increase in size rather than number to adapt to increasing workload demands (Karbassi et al., 2020; Zhao, Ye, Su, & Garg, 2020). Despite the large timeline that corresponds to a developmental transition from late infancy to early adulthood in humans, the number of branches holding the myofibrillar networks within cardiomyocytes is stable at about 3 branches per 10 sarcomere lengths. These relatively low branch numbers are also characteristic of cardiac muscles in healthy humans aged between 62 and 81 years (Z. Vue et al., 2023). In contrast to skeletal muscle, cardiomyocytes respond to aging by maintaining their sarcomere branching frequency.

The data herein suggests that the structural response to aging is muscle-type-specific. Overall, the present study underscores the importance of considering the ultrastructural changes within individual muscles when studying the effects of aging on normal physiology. Further research is warranted to elucidate the underlying mechanisms driving these observed structural changes over time and their functional consequences in the context of aging.

## Notes

### Competing Interest Statement

The authors have declared no competing interest.

## References

Ajayi, P. T., Katti, P., Zhang, Y., Willingham, T. B., Sun, Y., Bleck, C. K. E., & Glancy, B. (2022). Regulation of the evolutionarily conserved muscle myofibrillar matrix by cell type dependent and independent mechanisms. Nat Commun, 13(1), 2661. doi:10.1038/s41467-022-30401-9

Alway, S. E., Myers, M. J., & Mohamed, J. S. (2014). Regulation of satellite cell function in sarcopenia. Front Aging Neurosci, 6, 246. doi:10.3389/fnagi.2014.00246

Coggan, A. R., Spina, R. J., King, D. S., Rogers, M. A., Brown, M., Nemeth, P. M., & Holloszy, J. O. (1992). Histochemical and enzymatic comparison of the gastrocnemius muscle of young and elderly men and women. J Gerontol, 47(3), B71–76.

Dupont-Versteegden, E. E. (2005). Apoptosis in muscle atrophy: relevance to sarcopenia. Exp Gerontol, 40(6), 473–481. doi:10.1016/j.exger.2005.04.003

Goates, S., Du, K., Arensberg, M. B., Gaillard, T., Guralnik, J., & Pereira, S. L. (2019). Economic Impact of Hospitalizations in US Adults with Sarcopenia. J Frailty Aging, 8(2), 93–99. doi:10.14283/jfa.2019.10

Goldspink, G. (1970). The proliferation of myofibrils during muscle fibre growth. J Cell Sci, 6(2), 593–603.

Hinks, A., & Power, G. A. (2024). Age-related differences in the loss and recovery of serial sarcomere number following disuse atrophy in rats. bioRxiv, 2024.2006.2010.598222. doi:10.1101/2024.06.10.598222

Jorgenson, K. W., Hibbert, J. E., Sayed, R. K. A., Lange, A. N., Godwin, J. S., Mesquita, P. H. C., … Hornberger, T. A. (2024). A novel imaging method (FIM-ID) reveals that myofibrillogenesis plays a major role in the mechanically induced growth of skeletal muscle. eLife, 12. doi:10.7554/eLife.92674

Kamel, H. K., Maas, D., & Duthie, E. H., Jr. (2002). Role of hormones in the pathogenesis and management of sarcopenia. Drugs Aging, 19(11), 865–877. doi:10.2165/00002512-200219110-00004

Kaminska, A., Fidzianska, A., Schulze, G., Coper, H., Ossowska, K., Wolfarth, S., & Hausmanowa-Petrusewicz, I. (1998). Ultrastructural changes in the skeletal muscle of senile rats with significant age-dependent motor deficits. BASIC AND APPLIED MYOLOGY, 8(3), 185–190.

Karbassi, E., Fenix, A., Marchiano, S., Muraoka, N., Nakamura, K., Yang, X., & Murry, C. E. (2020). Cardiomyocyte maturation: advances in knowledge and implications for regenerative medicine. Nat Rev Cardiol, 17(6), 341–359. doi:10.1038/s41569-019-0331-x

Kim, Y., Ajayi, P. T., Bleck, C. K. E., & Glancy, B. (2022). Three-dimensional remodelling of the cellular energy distribution system during postnatal heart development. Philos Trans R Soc Lond B Biol Sci, 377(1864), 20210322. doi:10.1098/rstb.2021.0322

Lam, J., Katti, P., Biete, M., Mungai, M., AshShareef, S., Neikirk, K., … Hinton, A., Jr. (2021). A Universal Approach to Analyzing Transmission Electron Microscopy with ImageJ. Cells, 10(9). doi:10.3390/cells10092177

Lang, T., Streeper, T., Cawthon, P., Baldwin, K., Taaffe, D. R., & Harris, T. (2010). Sarcopenia: etiology, clinical consequences, intervention, and assessment. Osteoporosis international, 21, 543–559.

Larsson, L. (1978). Morphological and functional characteristics of the ageing skeletal muscle in man. A cross-sectional study. Acta Physiol Scand Suppl, 457, 1–36.

Larsson, L., Degens, H., Li, M., Salviati, L., Lee, Y. I., Thompson, W., … Sandri, M. (2019). Sarcopenia: aging-related loss of muscle mass and function. Physiological reviews, 99(1), 427–511.

Lin, I. H., Chang, J. L., Hua, K., Huang, W. C., Hsu, M. T., & Chen, Y. F. (2018). Skeletal muscle in aged mice reveals extensive transformation of muscle gene expression. BMC Genet, 19(1), 55. doi:10.1186/s12863-018-0660-5

Marzetti, E., & Leeuwenburgh, C. (2006). Skeletal muscle apoptosis, sarcopenia and frailty at old age. Exp Gerontol, 41(12), 1234–1238. doi:10.1016/j.exger.2006.08.011

Meltzer, H. Y., Kuncl, R. W., & Yang, V. (1976). Incidence of Z band streaming and myofibrillar disruptions in skeletal muscle from healthy young people. Neurology, 26(9), 853–857. doi:10.1212/wnl.26.9.853

Narici, M. V., & Maganaris, C. N. (2007). Plasticity of the muscle-tendon complex with disuse and aging. Exerc Sport Sci Rev, 35(3), 126–134. doi:10.1097/jes.0b013e3180a030ec

Oertel, G. (1986). Changes in human skeletal muscles due to ageing. Histological and histochemical observations on autopsy material. Acta Neuropathol, 69(3-4), 309–313. doi:10.1007/BF00688309

Pan, L., Xie, W., Fu, X., Lu, W., Jin, H., Lai, J., … Xiao, W. (2021). Inflammation and sarcopenia: A focus on circulating inflammatory cytokines. Exp Gerontol, 154, 111544. doi:10.1016/j.exger.2021.111544

Proctor, D. N., Balagopal, P., & Nair, K. S. (1998). Age-related sarcopenia in humans is associated with reduced synthetic rates of specific muscle proteins. J Nutr, 128(2 Suppl), 351S–355S. doi:10.1093/jn/128.2.351S

Roth, S. M., Martel, G. F., Ivey, F. M., Lemmer, J. T., Metter, E. J., Hurley, B. F., & Rogers, M. A. (2000). High-volume, heavy-resistance strength training and muscle damage in young and older women. J Appl Physiol (1985), 88(3), 1112–1118. doi:10.1152/jappl.2000.88.3.1112

Sayed, R. K. A., Hibbert, J. E., Jorgenson, K. W., & Hornberger, T. A. (2023). The Structural Adaptations That Mediate Disuse-Induced Atrophy of Skeletal Muscle. Cells, 12(24). doi:10.3390/cells12242811

Street, S. F. (1983). Lateral transmission of tension in frog myofibers: a myofibrillar network and transverse cytoskeletal connections are possible transmitters. J Cell Physiol, 114(3), 346–364. doi:10.1002/jcp.1041140314

Tabary, J. C., Tardieu, C., Tardieu, G., Tabary, C., & Gagnard, L. (1976). Functional adaptation of sarcomere number of normal cat muscle. J Physiol (Paris), 72(3), 277–291.

Thom, J. M., Morse, C. I., Birch, K. M., & Narici, M. V. (2007). Influence of muscle architecture on the torque and power-velocity characteristics of young and elderly men. Eur J Appl Physiol, 100(5), 613–619. doi:10.1007/s00421-007-0481-0

Visser, M., Goodpaster, B. H., Kritchevsky, S. B., Newman, A. B., Nevitt, M., Rubin, S. M., … Harris, T. B. (2005). Muscle mass, muscle strength, and muscle fat infiltration as predictors of incident mobility limitations in well-functioning older persons. J Gerontol A Biol Sci Med Sci, 60(3), 324–333. doi:10.1093/gerona/60.3.324

Vue, Z., Ajayi, P. T., Neikirk, K., Murphy, A. C., Prasad, P., Jenkins, B. C., … Hinton, A., Jr. (2023). Human Heart Failure Alters Mitochondria and Fiber 3D Structure Triggering Metabolic Shifts. bioRxiv. doi:10.1101/2023.11.28.569095

Vue, Z., Garza-Lopez, E., Neikirk, K., Katti, P., Vang, L., Beasley, H., … Murphy, A. C. (2023). 3D reconstruction of murine mitochondria reveals changes in structure during aging linked to the MICOS complex. Aging Cell, 22(12), e14009.

Williams, P. E., & Goldspink, G. (1976). The effect of denervation and dystrophy on the adaptation of sarcomere number to the functional length of the muscle in young and adult mice. J Anat, 122(Pt 2), 455-465.

Willingham, T. B., Kim, Y., Lindberg, E., Bleck, C. K. E., & Glancy, B. (2020). The unified myofibrillar matrix for force generation in muscle. Nature Communications, 11(1). doi:10.1038/s41467-020-17579-6

Yamada, M., Kimura, Y., Ishiyama, D., Otobe, Y., Suzuki, M., Koyama, S., & Arai, H. (2022). Combined effect of lower muscle quality and quantity on incident falls and fall-related fractures in community-dwelling older adults: A 3-year follow-up study. Bone, 162, 116474. doi:10.1016/j.bone.2022.116474

Yu, J. G., Carlsson, L., & Thornell, L. E. (2004). Evidence for myofibril remodeling as opposed to myofibril damage in human muscles with DOMS: an ultrastructural and immunoelectron microscopic study. Histochem Cell Biol, 121(3), 219–227. doi:10.1007/s00418-004-0625-9

Zhao, M. T., Ye, S., Su, J., & Garg, V. (2020). Cardiomyocyte Proliferation and Maturation: Two Sides of the Same Coin for Heart Regeneration. Front Cell Dev Biol, 8, 594226. doi:10.3389/fcell.2020.594226

